# On the expression of co-operative feeding behaviour in 3^rd^ instar *Drosophila melanogaster* larvae!

**DOI:** 10.1101/678631

**Authors:** Lucas Khodaei, Tara Newman, Samantha Lum, Henry Ngo, Matthew Maoloni, Tristan A.F. Long

## Abstract

Under poor nutritional conditions, 3^rd^ instar *Drosophila melanogaster* larvae will work collaboratively in feeding clusters to obtain resources that cannot be reached individually. To better understand the conditions that influence the expression of this behaviour we examined the frequencies, the size and the membership in vials of flies that were initially seeded with either 100 or 200 eggs each using flies from both a large, outbred population and a replicate population that was homozygous for the *bw* allele. Overall, more feeding clusters, containing more larval participants were observed in the higher density vials compared to the lower density vials, consistent with the idea that this social behaviour is a response to dwindling resources in the environment. The presence of the *bw* allele did not result in greater egg-to-adult mortality, nor did it result in lower participation in feeding clusters.

## 1. Introduction

The fruit fly, *Drosophila melanogaster*, has a rich and celebrated history as a model organism in scientific research, and whose study has contributed to some of the greatest biological discoveries of all time (Weiner 1999, Brookes 2001). In recent years, it has also become a popular model species in which the ultimate and proximate causes of social behaviour can be examined (*e.g.* Wu *et al.* 2003, Sokolowski 2010, Durisko & Dukas 2013, Durisko *et al*. 2014, Camiletti & Thompson 2016).

Of the many interesting behaviours exhibited by *D. melanogaster*, our study focuses on the expression of social feeding among 3^rd^ instar larvae (*see* Dombrovski *et al*. 2017). In this species, females lay eggs in fermenting vegetation (Reaume & Sokolowski 2006), where their offspring are often confined until their eclosion as adults (Sokolowski 1985). The structural and nutritional environment experienced by larval *D. melanogaster* often deteriorates over time, through the combined action of their burrowing and feeding, the buildup of their waste products, the effects of secreted digestive enzymes, and the activity of the microbial community. While desirable resources may potentially be found at lower stratums, the liquidification of the upper layers of the larvaes’ habitat makes it difficult for them to access them, as they risk suffocation (Dombrovski *et al*. 2017). In these situations, individuals aggregate to form “feeding clusters” by synchronizing their digging movements to dig a ‘well’ (*sensu* Louis & de Polavieja 2017) that allows them to gain access to deeply buried resources, while still allowing respiration (using their posterior breathing spiracles) via a common air cavity (Wu *et al*. 2003, Justice *et al*. 2012, Dombrovski *et al*. 2017).

This study was undertaken to better understand some of the factors that may potentially influence the expression of this group feeding behaviour, as part of a larger investigation of this social phenomenon (Khodaei & Long 2019). Specifically, we set out to examine whether this phenomenon was influenced by larval density, and/or the genetic background of the larvae.

First, we explored the extent to which the formation and appearance of feeding clusters is associated with the local density of larvae. The nutritional environment experienced by larvae is often influenced by the abundance of conspecifics, with higher densities associated with greater competition for resources, slower development, higher mortality, and worsened individual condition (*reviewed in* Ashburner *et al*. 2005, Tennant *et al*. 2014). Since the local density of larvae is likely to be strongly associated with the rate/amount of resource depletion and/or environmental degradation, it is possible that larvae will be more likely to engage in group foraging at higher densities in order to access needed resources. However, as *D. melanogaster* larval cannibalism is expressed in times of nutritional stress and can involve individuals consuming other larvae (Vijendravarma *et al*. 2013), adult carcasses (Yang 2018) and/or eggs (Ahmad *et al*. 2015; *but see* Narasimha *et al*. 2019), at higher densities one might alternatively see less social behaviour being expressed, if individuals become more adverse to being in close contact with hungry conspecifics.

In their study of group feeding, Dombrovski *et al*. (2017) found that both mechanosensory cues and vision are critical for the efficient co-ordination of feeding behaviours between adjacent larvae. The importance of sight was demonstrated in one of their assays as larvae from three different vision-impaired mutant lines, norpA^P41^, GMR-hid1, and GMR-hid2, formed fewer, and smaller feeding clusters than flies from a wild-type population. In Khodaei & Long (2019), our focal larvae were obtained from a *wild-type* population of *D. melanogaster*, but competed in some of our assays with flies from a replicate population where individuals homozygous for the *bw* allele (where adults have brown coloured eyes). While adult *bw* homozygotes may have have compromised vision that renders them selectively inferior to *wild-type* individuals (*e.g*. MacLellan *et al*. 2009, but see Burnet *et al*. 1968), whether there are any phenotypic differences with respect to larval group foraging is unknown.

## 2. Materials and methods

For our assays we used *Drosophila melanogaster* that we obtained from two populations: *Ives* (hereafter ‘IV‘), and IV-*bw*. The IV population is an outbred, *wild-type* population which originated from a sample of 200 mated females collected in Amherst MA, USA in 1975 (Rose 1984). The IV-*bw* population was created through the introgression of the recessive, brown-eyed allele, *bw* into the IV genetic background via repeated back-crossing. Both of these populations are kept at large sizes (∼3500 adults per generation) in non-overlapping generations, and are maintained under standardized conditions (incubated at 25°C, 60% RH on a 12hr-L:12hr-D diurnal light cycle) (Martin & Long 2015). Flies are cultured in narrow *Drosophila* vials containing 10mL of a banana, killed-yeast and agar media, set at an initial density of approximately of 100 eggs/vial. Populations are cultured *en masse* every 14 days (Rose 1984, Martin & Long 2015), by mixing all eclosed adult flies together under light CO_2_ anesthetic, distributing them into vials containing fresh media and a small amount of live yeast for 2-3 hours before being removed. The eggs laid in these vials are then culled by hand to the appropriate density, and these vials are used to found the next generation of culture.

In our experiment we set out to examine survivorship and co-operative behaviour of both the IV and IV-*bw* populations at two initial egg density treatments: 100 eggs/vial (the typical culture density), as well as 200 eggs/vial. Under these more crowded conditions, IV flies may experience more resource limitation (Tennant *et al*. 2014). We established 26 replicate vials for each of these four experimental treatments. Eggs were obtained by placing adult IV and IV-*bw* flies into oviposition chambers that contained grape juice/agar medium (Sullivan *et al*., 2000) for ∼18h (overnight). From these surfaces we carefully counted and transferred sets of 100 or 200 eggs into new vials containing 10ml of standard media before being placed in the incubator.

Starting 24h after the experimental vials’ creation, we began surveying them for signs of larval cooperative feeding. Surveys were done on 3 consecutive days in 3 sessions/day, spaced 3h apart by a team of well-trained observers who were blind to the treatment identity of the vials. These observers counted the total number of feeding clusters present in each vial, where a cluster was defined as a grouping of 3 or more larvae which were all feeding in a downward direction creating a depression in the food media. Feeding clusters were defined as being separate from one another if there was evidence that larvae in the two groups were not working together (*i.e*. oriented in opposite directions) and were separated from each other by at least 0.5mm. For those clusters that had formed along the side of the vial, we measured the width (the distance from one side of the foraging group to the other at the food surface level) and depth (the distance from the food surface level to the lowest point in the foraging group) of each of the clusters using a caliper to the nearest 0.5 mm. We estimated the size of each of the cluster‘s depression observed as the area of a triangle using our width and depth measurements, and then calculated the mean cluster depression areas measured across all sessions. The total number of larvae along the contour of each feeding cluster that were visible through the transparent wall was also counted.

Fourteen days after the experimental vials’ creation, all eclosed adult flies were carefully removed from the vial using light CO_2_ anesthesia and were counted.

All statistical analyses were performed using R version 3. 3.2 (R Core Team 2016) with vials as the unit of replication. We compared the egg-to-adult survivorship rates using a generalized linear model (GLM), with a quasibinomial error distribution, where the response variable was the number of flies (out of the initial number of eggs in the vial) that survived to adulthood, and the independent variables were the population source, the initial egg density and their interaction. We determined the statistical significance of independent variables using a likelihood-ratio Chi-square test using the *Anova* function in the *car* package (Fox & Weisberg 2011).

We compared the frequency of cluster formation using a GLM with quasipoisson error distribution, where the independent variables were the population source, the initial egg density and their interaction, and the dependent variable was the number of clusters in each vial summed across all observation sessions. As above, we used a likelihood-ratio Chi-square test to determine the statistical significance of independent variables.

As both the data on mean cluster size and mean number of larvae present met parametric assumptions, we examined the effects of population origin, initial egg density and their interaction using the Two-Way Analysis of Variance (ANOVA) method, which was followed by a *post-hoc* Tukey HSD test to compare the means of the treatments, if necessary.

## 3. Results

When comparing the rates of egg-to-adult survivorship across our experimental treatments (Figure 2), we found a significant effect of density (LR χ^2^=172.57, df=1, p<0.0001), but no difference between the IV and IV-populations (LR χ^2^= 1.29, df=1, p=0.25) and no significant interaction between population and density (LR χ^2^=0.67, df=1, p=0.42).

**Figure 1:**
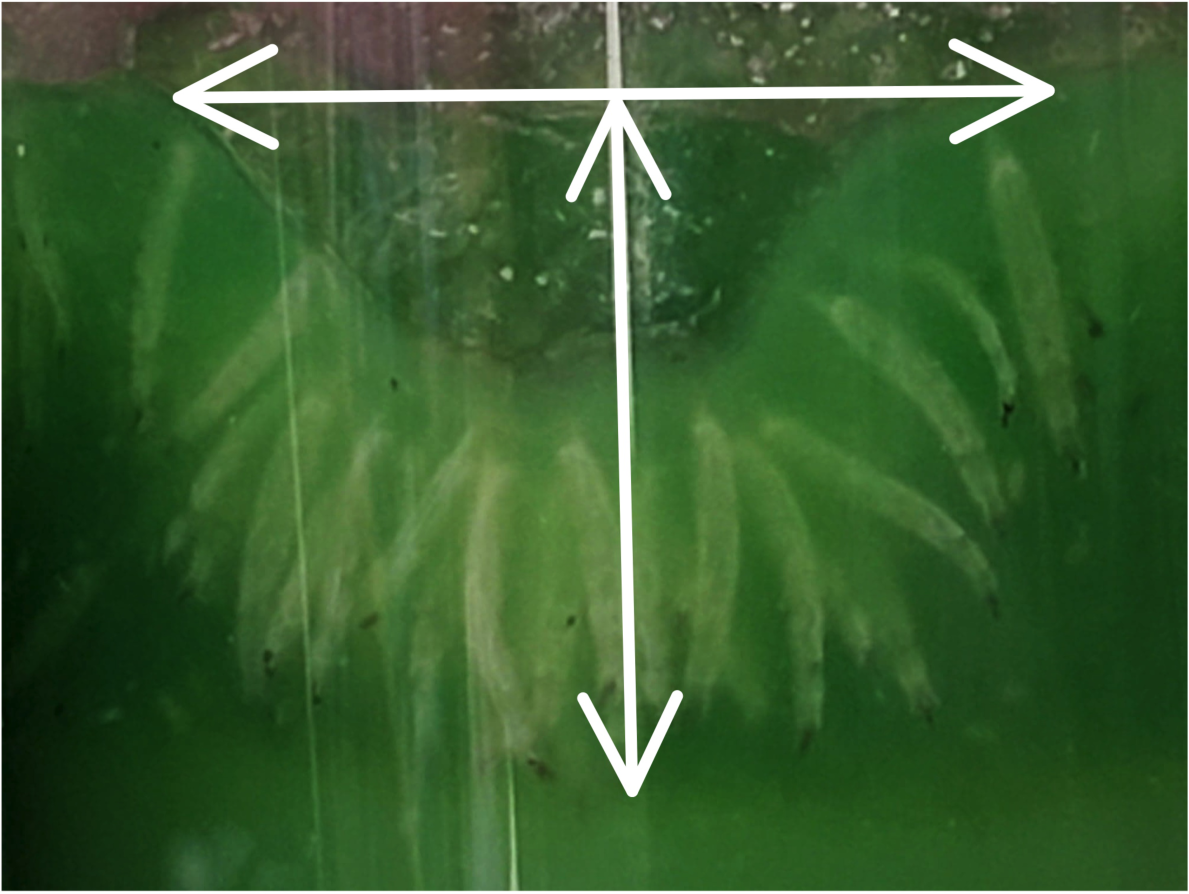
Photo depicting a *Drosophila melanogaster* larval feeding cluster with lines illustrating how width and depth of each cluster were measured. The width of each cluster was defined as the horizontal distance between the outermost cluster group members, while the depth of each cluster was defined as the vertical distance from the media surface level to the lowest point in the foraging group. We also counted the number of larvae along the contour of the feeding cluster that were visible through the transparent wall of the vial as an index of how many larvae were co-operating.

**Figure 2:**
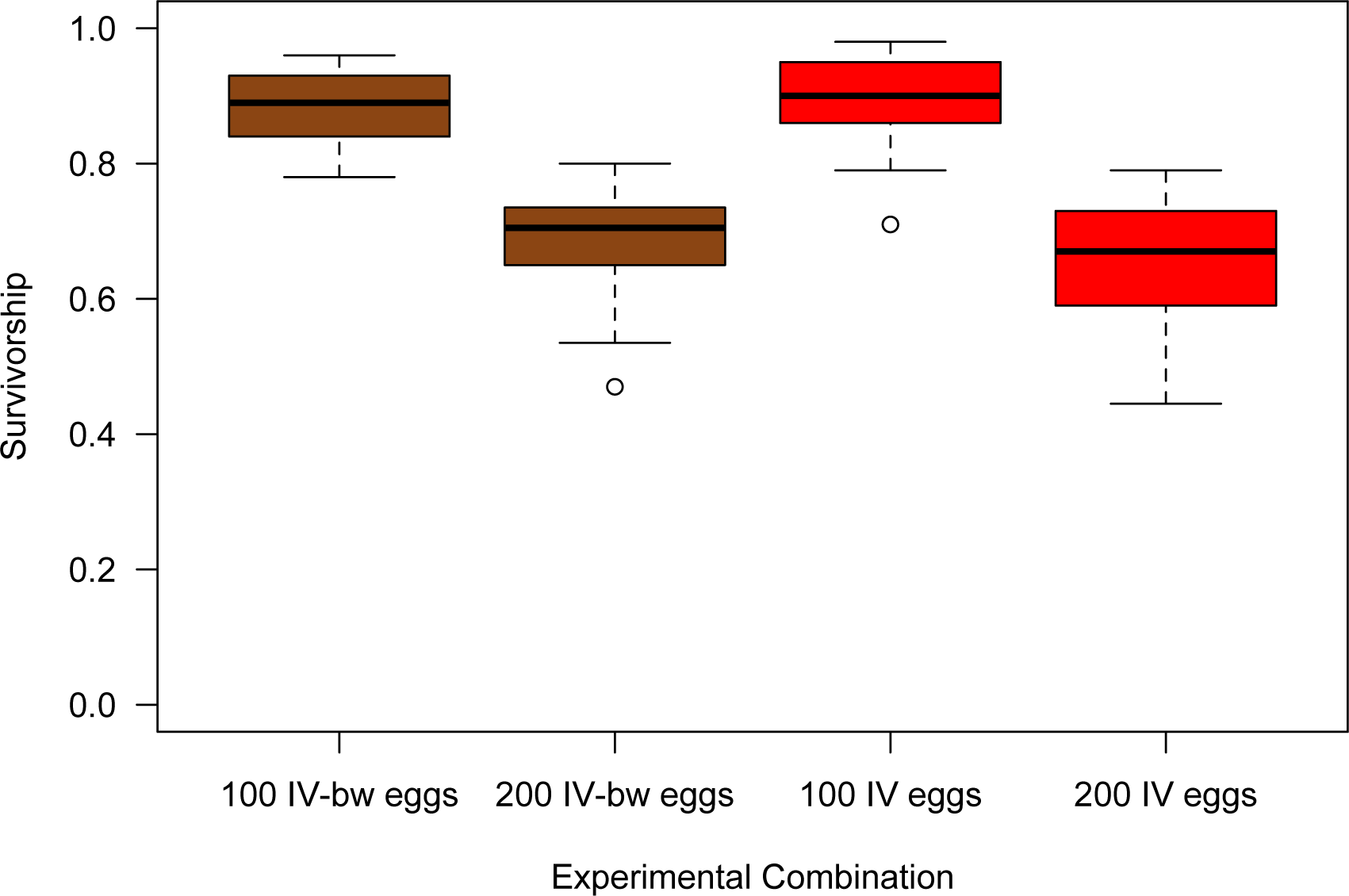
Boxplots illustrating egg-to-adult survivorship rates in vials of *Drosophila melanogaster* obtained from the IV or the IV-*bw* that were initiated with either 100 or 200 eggs. Each box encloses the middle 50% of data (Inter-Quartile Range, IQR), with the location of the median represented by a horizontal line. Values >±1.5 x the IQR outside the box are considered outliers and depicted as open circles. Whiskers extend to the largest and smallest values that are not outliers.

The total number of feeding clusters (Figure 3) observed was also greater in vials that had been seeded with 200 eggs compared to those from the 100 egg/vial treatment (LR χ^2^=11.09, df=1, p=0.0009). We found no significant difference between the vials that contained IV larvae and those which contained IV-*bw* larvae (LR χ^2^= 0.06, df=1, p=0.8), and no significant interaction between the population origin and the egg density treatments (LR χ^2^=0.21, df=1, p=0.65).

**Figure 3:**
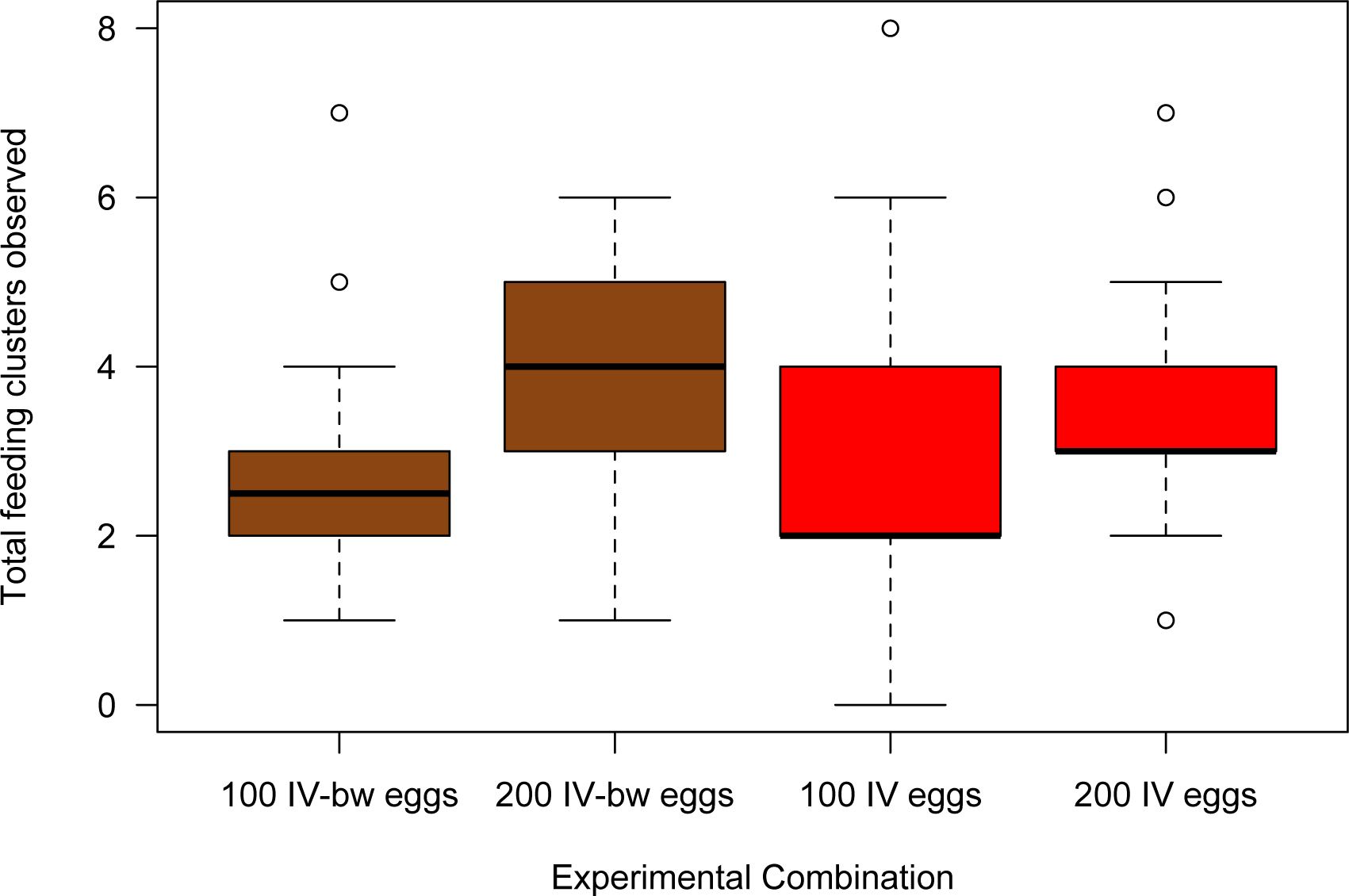
Boxplots illustrating the total number of larval feeding clusters observed in vials of *Drosophila melanogaster* obtained from the IV or the IV-*bw* population that were initiated with either 100 or 200 eggs. Boxplot components as described as in Figure 2.

When examining variation in the size of feeding clusters (Figure 4), we found no significant effect of either population origin (F=1.73, df=1,99, p=0.19), egg density (F=0.01, df=1,99, p=0.91) or their interaction (F=2.70, df=1,99, p=0.10).

**Figure 4.**
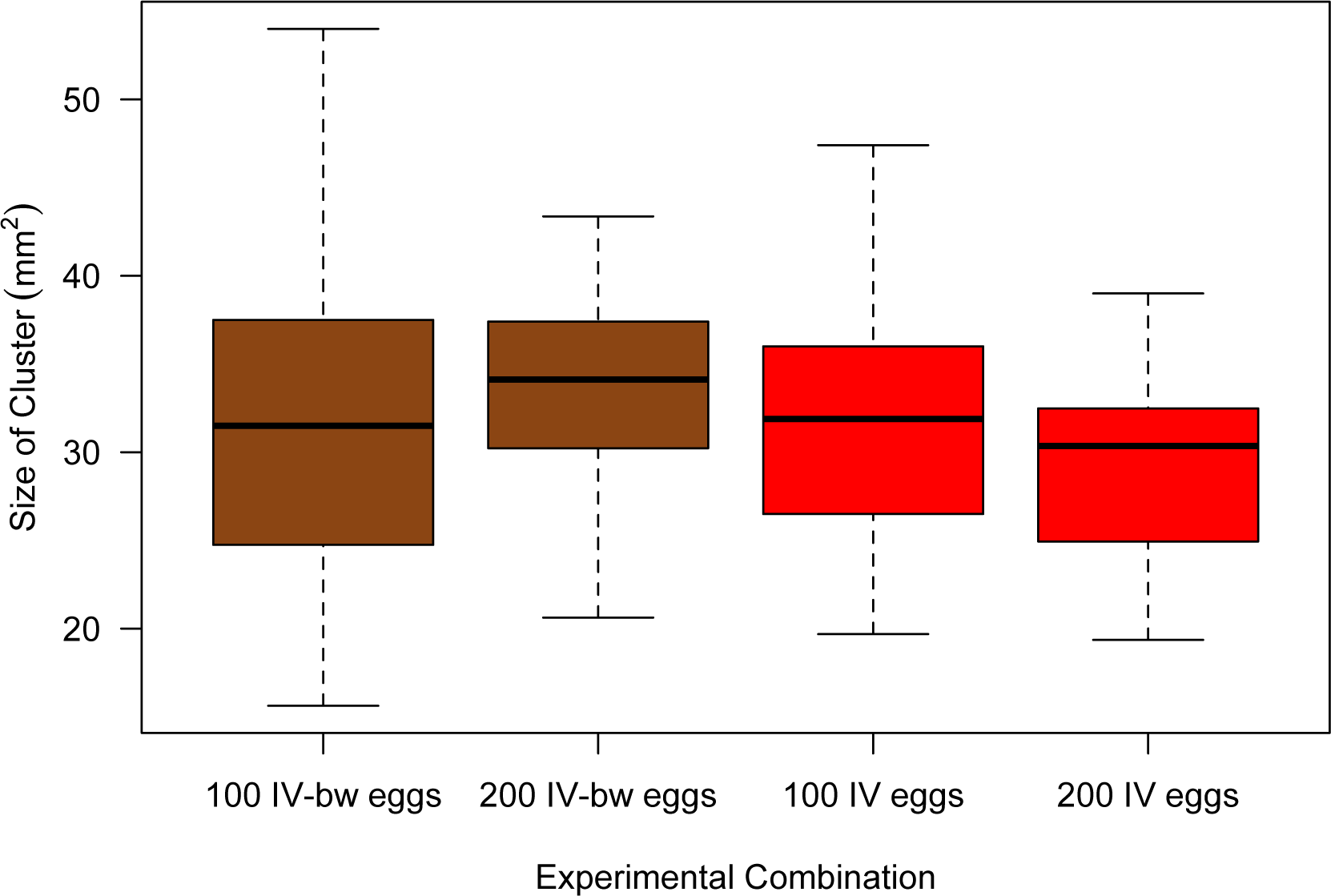
Boxplots illustrating the mean size of all larval feeding clusters observed in vials of *Drosophila melanogaster* obtained from the IV and IV-*bw* populations that were initiated with either 100 or 200 eggs. Boxplot components as described as in Figure 2.

Finally, when we examined the number of larvae found in each cluster (Figure 5) the main effect of population origin was not significant (F=0.70, df=1,99, p=0.40). There were, however, more larvae on average in clusters in those vials that had been seeded with 200 eggs compared to those from the 100 egg/vial treatment (F=14.85, df=1,99, p=0.0002) and there was a significant interaction between the two main factors (F=11.15, df=1,99, p=0.001). Specifically, we observed significantly more larvae, on average, in clusters from the IV-*bw* 200 treatment vials, compared to the three other treatment combinations 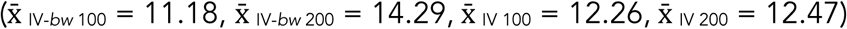.

**Figure 5.**
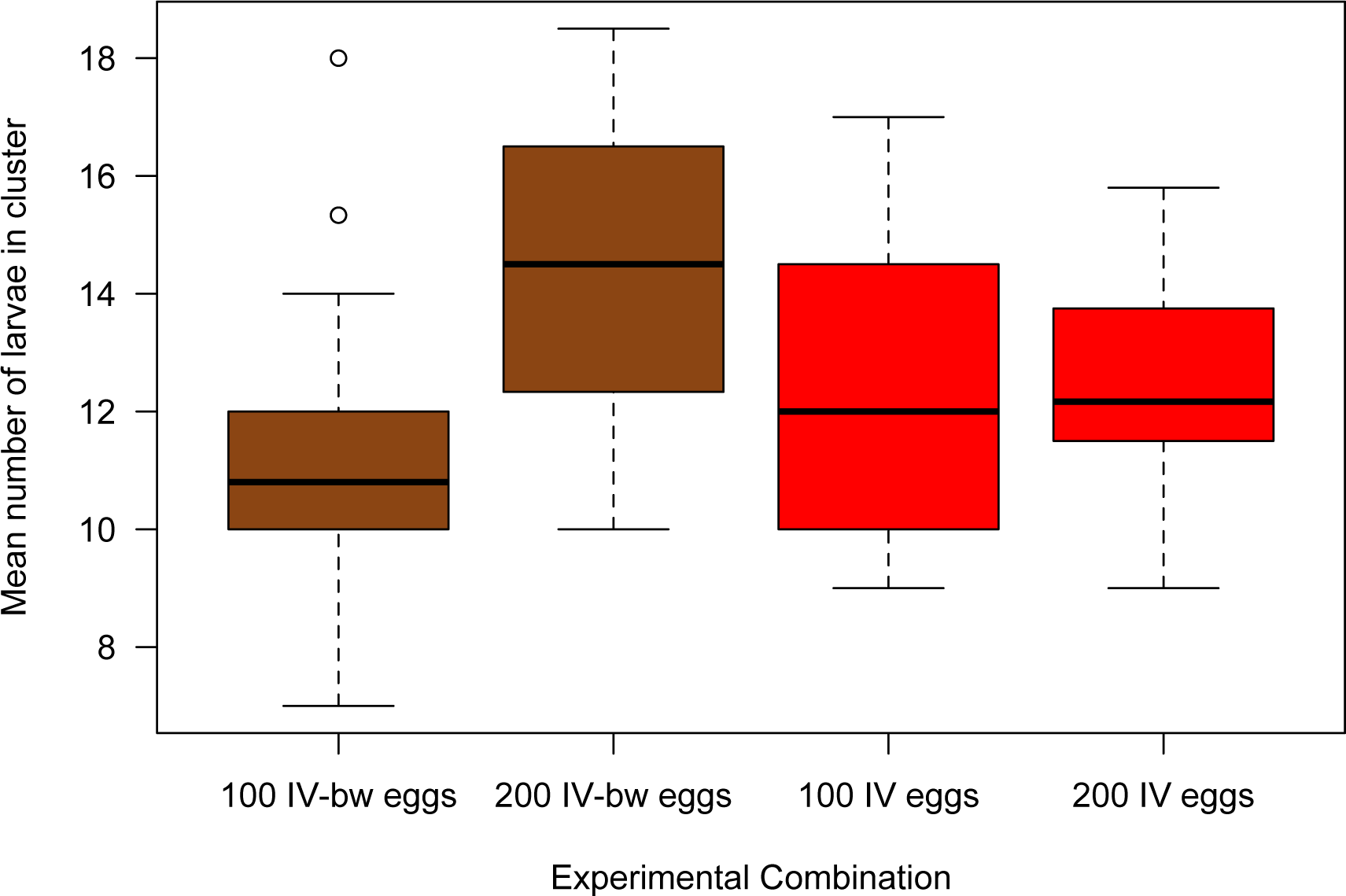
Boxplots illustrating the mean number of larvae present in all larval feeding clusters observed in vials of *Drosophila melanogaster* obtained from the IV and IV-*bw* populations that were initiated with either 100 or 200 eggs. Boxplot components as described in Figure 2.

## 4. Discussion

In this study we set out to examine whether the manifestation of collaborative feeding clusters in 3^rd^ instar *D. melanogaster* larvae was influenced by the local density of conspecifics in their environment and/or whether there were differences between larvae obtained from two populations that differed at the *bw* locus. Overall, we saw that in the more resource-depleted high-density vials, that co-operative feeding clusters were more commonly observed. Furthermore, we saw no evidence that larvae homozygous for *bw* allele were less likely to participate in group feeding behaviours.

Increasing the density of eggs in the vials clearly led to an environment that was more challenging to the development of our *D. melanogaster* larvae, which was reflected in our lower egg-to-adult survival rates in those vials that had been initially seeded with 200 eggs compared to those that had initially contained 100 eggs apiece (Figure 2). This is consistent with many other studies that have documented the adverse effects associated with the more inhospitable environment that results from increased conspecific densities (*reviewed in* Ashburner *et al*. 2005, Tennant *et al*. 2014). Successful larval development in *D. melanogaster* requires individuals to ingest between 3 to 5 times their body weight in yeasts and decaying matter each day (Chiang & Hodson 1950), and consequently resources will become depleted more quickly as the number of developing larvae within a vial increases. Thus, seeing more feeding clusters within the high-density vials (Figure 3) is indicative that the cooperative feeding behaviour is dependent on the quality of the larvae‘s environment. The burrowing action of larvae causes the upper layers of their media to become both nutrient-poor, and increasingly liquefied (Dombrovski *et al*. 2017), which would only be exacerbated by larger numbers of conspecifics.

Despite observing more clusters in higher density vials, we did not see any statistically significant difference in the size of the feeding clusters compared to the lower density vials (Figure 4). This somewhat surprising negative result suggests several possibilities, which may be worth pursuing in future studies. First, there may be an optimal size for feeding clusters, where individuals obtain the greatest resources in return for their efforts, and that if clusters grow beyond that point, individuals may move to smaller clusters with greater net benefits. Secondly, the upper layers of media in the higher density vials may be more fluid than those in the lower density vials, and whose structural instability may prevent larvae in the 200 egg vials from successfully excavating deeper wells. Finally, the 3^rd^ instar larvae in the higher density vials may be of worse physiological condition (due to more limited nutritional resources) and may be less capable of the sustained, co-ordination of their movements needed to create a larger air cavity.

When comparing the number of larvae within each cluster (Figure 5) we observed that, on average, feeding clusters in the higher density vials had ∼1.67 more members than those from the lower density vials (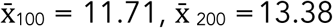; 2 sample t-test, t = −3.66, df = 101, p = 0.0004), which could reflect the greater motivation to work together in the more challenging environment and/or the greater absolute number of larvae present in the higher density vials. However, the difference between 100 and 200 egg vials was much larger in those vials that contained IV-*bw* larvae than those that contained IV larvae, which resulted in a significant population or origin x density treatment interaction. It is unclear what may have contributed to this observed difference. Apart from this one situation, we did not see any other evidence that larvae from the IV-*bw* population differed in their larval feeding behaviour, or response to differences in larval density from those larvae obtained from the IV population. If being homozygous at the *bw* allele impaired the larvae‘s’ ability to work collectively, we would have expected to see fewer clusters, smaller clusters and/or fewer larvae in clusters when larvae originated from the IV-*bw* population. None of these predicted differences were observed. While the phenotype associated with *bw* homozygosity can be deleterious to adults (MacLellan *et al*. 2009, Colpitts *et al*. 2017), in our study, it does not appear to interfere with their ability to feed collectively as larvae (Figures 3, 4, & 5), nor have a deleterious effect on egg-to-adult survivorship (Figure 1).

This study contributes to the growing body of research examining the proximate and ultimate causes of group feeding behaviour in *Drosophila melanogaster* (Wu *et al*. 2003, Justice *et al*. 2012, Dombrovski *et al*. 2017, Louis & de Polavieja 2017), and provides insight into the conditions that favour the evolution of social behaviours.

## Supporting information

Surivorship Data

Cluster Data

## Data Accessibility Statement

Please see the Supplemental Cluster Data and Supplemental Survivorship Data files associated with this manuscript.

## Author Contributions

The project was conceived by TAFL and LK. Counting of eggs, clusters, and/or adult flies was performed by LK, TN, SL, DH, MM and/or TAFL. TAFL wrote the manuscript.

## Acknowledgements

TAFL was funded with a Natural Sciences and Engineering Research Council (NSERC) Discovery grant. We would like to thank the anonymous referee who was the inspiration for these experiments. This work was conducted at Wilfrid Laurier University, which exists on the traditional territory of the Neutral, Anishnawbe, and Haudenosaunee peoples.

